# Alpha Herpesvirus Egress and Spread from Neurons Uses Constitutive Secretory Mechanisms Independent of Neuronal Firing Activity

**DOI:** 10.1101/729830

**Authors:** Anthony E. Ambrosini, Nikhil Deshmukh, Michael J. Berry, Lynn W. Enquist, Ian B. Hogue

**Affiliations:** Department of Molecular Biology, and Princeton Neuroscience Institute, Princeton University, Princeton, NJ USA; Center for Immunotherapy, Vaccines, and Virotherapy, Biodesign Institute, and School of Life Sciences, Arizona State University, Tempe, AZ USA

## Abstract

Alpha herpesviruses naturally infect the peripheral nervous system, and can spread to the central nervous system causing severe deadly or debilitating disease. Because alpha herpesviruses spread along synaptic circuits, and infected neurons exhibit altered electrophysiology and increased spontaneous firing, we hypothesized that alpha herpesviruses use activity-dependent synaptic vesicle-like regulated secretory mechanisms for egress and spread from neurons. To address this hypothesis, we used a compartmentalized primary neuron culture system to measure egress and spread of pseudorabies virus (PRV), pharmacological and optogenetics approaches to modulate neuronal firing activity, and a live-cell fluorescence microscopy assay to directly visualize the exocytosis of individual virus particles from infected neurons. Using tetrodotoxin to silence neuronal activity, we observed no inhibition of virus spread, and using potassium chloride or optogenetics to elevate neuronal activity, we also show no increase in virus spread. Using a live-cell fluorescence microscopy method to directly measure virus egress from infected neurons, we observed no association between virus particle exocytosis and intracellular Ca^2+^ signaling. Finally, we observed virus particle exocytosis occurs in association with constitutive secretory Rab GTPases, Rab6a and Rab8a, not Rab proteins that are associated with the Ca^2+^-regulated secretory pathway in neurons, Rab3a and Rab11a. Therefore, we conclude that alpha herpesvirus egress and spread is independent of neuronal activity and Ca^2+^ signaling because virus particle exocytosis uses constitutive secretory mechanisms in neurons.

**Author Summary:** Alpha herpesviruses, including important human pathogens Herpes Simplex Virus 1 and 2, and Varicella-Zoster Virus, are among the very few viruses that naturally infect the nervous system. These viruses cause recurrent herpetic and zosteriform lesions, peripheral neuropathies, and deadly or debilitating central nervous system diseases. Many of the molecular and cellular mechanisms of viral egress and spread remain unknown, particularly in the context of specialized neuronal cell biology. Our results indicate that elevated firing activity of infected neurons is not functionally or mechanistically linked to virus egress and spread; therefore, therapies targeting peripheral neuropathic symptoms, elevated neuronal activity, and synaptic vesicle secretory mechanisms are unlikely to affect virus spread in the nervous system.

## Introduction

The alpha herpesviruses, including herpes simplex virus 1 and 2 (HSV-1 & -2), varicella-zoster virus (VZV), and pseudorabies virus (PRV; suid herpesvirus 1), are among the very few viruses that have evolved to productively exploit the mammalian nervous system. After initial infection of epithelial tissues, alpha herpesviruses efficiently enter and establish life-long latency in sensory and autonomic peripheral nervous system (PNS) neurons. Upon reactivation of viral replication, virus particles undergo anterograde axonal transport, exocytosis, and spread back to peripheral tissues, where they cause characteristic recurrent herpetic or zosteriform lesions. Rarely in natural hosts, but more frequently in young, old, immunocompromised, and non-natural host species, alpha herpesviruses can also spread into the central nervous system (CNS), causing acute neurological disease like herpes encephalitis [1]. It is increasingly apparent that alpha herpesviruses can also spread to the CNS asymptomatically, even in healthy, immunocompetent, natural hosts, which might contribute to the development of chronic neurodegenerative diseases [2].

In humans, HSV and VZV cause a variety of sensory neuropathies, including numbness, tingling, itch, and burning pain. In non-natural hosts (e.g. laboratory rodents and many other domesticated animals), PRV causes an intense pruritus known as “mad itch” [3]. These neuropathic symptoms are thought to be caused by altered electrophysiology of infected neurons. HSV is reported to alter neuron excitability by downregulating expression and cell-surface localization of voltage-gated and inward rectifying ion channels [4–9], yet PRV and some syncytial strains of HSV are also reported to induce synchronized spontaneous neuronal activity *in vitro* and in non-myelinated neurons *in vivo* [4–8]. Using PRV reverse genetics, the Enquist laboratory previously showed that two viral membrane proteins are required for elevated spontaneous activity of neurons *in vitro* and *in vivo*: 1) membrane glycoprotein gB is part of the core fusion complex that mediates membrane fusion during viral entry and cell-cell fusion leading to syncytia; 2) membrane protein US9 (together with glycoproteins gE and gI) is required for vesicular transport of PRV particles and glycoproteins into the axon of neurons. These findings lead to a model where viral membrane fusion proteins are transported into the axons of infected neurons, form fusion pores between adjacent non-myelinated axons, and cause the neurons to become electrically coupled. This leads to elevated and synchronous activity propagating through the network [10,11].

Constitutive secretory mechanisms mediate secretion of extracellular cargoes in all cell types, as well as homeostatic maintenance of lipids and proteins that form the plasma membrane itself. In general, protein cargoes are sorted at the trans-Golgi network (TGN) and recycling/sorting endosomes into discrete secretory vesicles. Rab-family GTPases bind the cytosolic face of intracellular membranes to specify organelle identity and regulate essentially all intracellular membrane trafficking. Rab6 and Rab8 proteins have been shown to associate with TGN and post-Golgi secretory vesicles to mediate their constitutive exocytosis [12–14]. Rab11 proteins function at the interface between endocytic and secretory pathways to regulate constitutive exocytosis of both post-Golgi secretory vesicles and recycling of endocytic cargoes to the plasma membrane [12,13]. Previously, we demonstrated that constitutive secretory Rab proteins, Rab6a, Rab8a, and Rab11a, are associated with exocytosis of PRV particles in non-neuronal cells [14,15], and the Elliott laboratory demonstrated that Rab6 and Rab11 are also required for HSV-1 glycoprotein trafficking and secondary envelopment [16,17].

In neurons and many other professional secretory cell types, exocytosis of signaling molecules such as neurotransmitters and hormones is highly regulated by intracellular Ca^2+^. Rab3 proteins are present on synaptic vesicles in neurons and are required for Ca^2+^-regulated exocytosis [18]. Rab11 is reported to play divergent roles in constitutive versus Ca^2+^-regulated exocytosis: Rab11 proteins participate in constitutive exocytosis in non-neuronal cells, as described above, but are reported to co-localize with Rab3 on synaptic vesicles on neurons, and mediate Ca^2+^-regulated, not constitutive, exocytosis in neuroendocrine cells [19]. Several reports in the alpha herpesvirus literature suggest an association between Ca^2+^-regulated secretory machinery and assembly/egress in specialized cell types. Rab3a and other synaptic vesicle markers were detected colocalized with HSV-1 proteins and virus particles in neurons [20], and Rab27a, which colocalizes with Rab3 on Ca^2+^-regulated secretory vesicles, was reported to be required for efficient HSV-1 replication in an oligodendrocyte cell line [21].

Since alpha herpesvirus infection causes elevated neuronal activity *in vitro* and *in vivo*, and viral proteins and particles are reported to be associated with synaptic vesicle/Ca^2+^-regulated secretory factors, we hypothesized: 1. PRV egress from neurons uses Ca^2+^-regulated exocytosis mechanisms; and 2. elevated firing activity would promote efficient viral egress and spread from neurons. To address these hypotheses, we took advantage of a modified Campenot tri-chamber neuronal culture system to measure transneuronal egress and spread of PRV, pharmacological and optogenetic approaches to modulate neuronal activity, and a live-cell fluorescence microscopy assay of virus particle exocytosis from primary neurons. Contrary to these hypotheses, we found that PRV egress and spread is not dependent on neuronal firing activity because PRV particle exocytosis uses constitutive secretory mechanisms, even in neurons.

## Results

### Tetrodotoxin and KCl affect firing activity of PRV-infected SCG neurons

In primary superior cervical ganglion (SCG) neurons *in vitro*, PRV causes a significant increase in firing activity by 8-10 hours post-infection (hpi) [10], and a significant increase in intracellular Ca^2+^ accumulation [22]. Tetrodotoxin (TTX) prevents neuronal firing by blocking voltage-gated Na^+^ channels, and potassium chloride (KCl) directly excites neurons by depolarizing their membrane potential. To ensure that TTX and KCl function in the context of PRV infection, we infected primary SCG cultures with PRV 468, expressing the fluorescent protein Ca^2+^ sensor GCaMP3, and monitored their Ca^2+^-dependent fluorescence between 9-11 (hpi), in the presence or absence of these drugs. TTX (4 μM) was added beginning at 4 hpi, and KCl (55 mM) was perfused in pulses beginning at 6 hpi. Untreated neurons exhibited an intermediate variation in Ca^2+^-dependent fluorescence over time (Fig 1A), and an intermediate steady-state accumulation of Ca^2+^-dependent fluorescence (Fig 1B). TTX treatment effectively silenced neuronal activity and reduced steady-state Ca^2+^ accumulation, whereas KCl treatment increased the variance and accumulation in Ca^2+^-dependent fluorescence over time (Fig 1).

**Fig 1.**
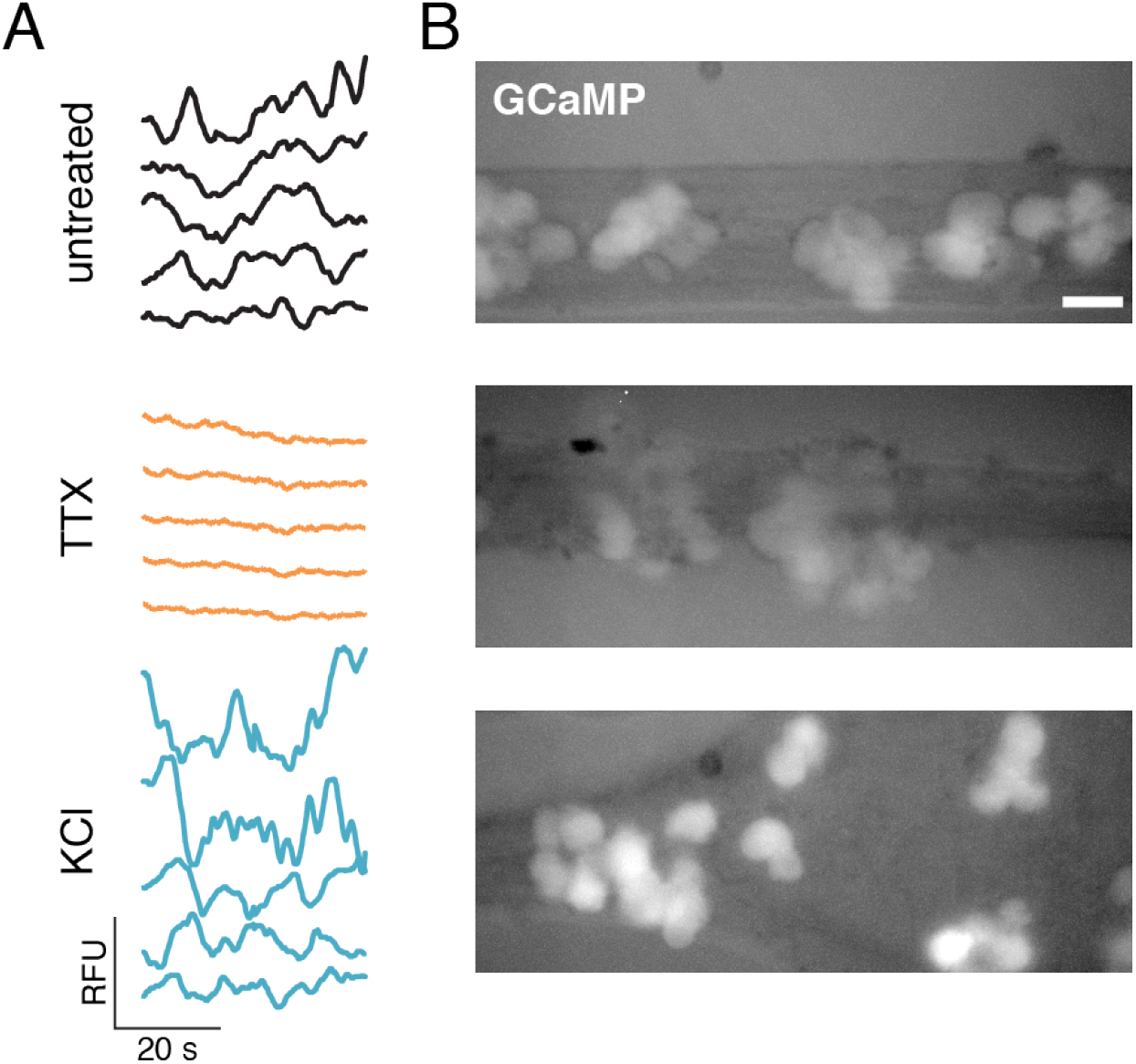
Tetrodotoxin and KCl affect activity of infected neurons. Primary SCG neurons were infected with a recombinant PRV expressing the fluorescent Ca^2+^ sensor GCaMP3 and treated with 4μM tetrodotoxin (TTX), pulses of 55mM KCl, or untreated, as indicated. (A) Relative Ca^2+^-dependent GCaMP3 fluorescence of 5 representative cells over time. (B) Steady-state Ca^2+^ accumulation indicated by GCaMP3 fluorescence. Scale bar represents 40 μm.

### Neuronal activity does not promote PRV spread from neurons to non-neuronal cells

To determine whether neuronal activity affects spread of PRV from SCG neurons to non-neuronal recipient cells, we took advantage of a modified Campenot tri-chamber culture system that fluidically separates SCG cell bodies from their axons [23]. Primary SCG neurons were seeded into the left soma chamber. After approximately 2 weeks in culture, SCG neurons extend axons under the chamber walls into the rightmost neurite chamber. After 3 weeks in vitro, non-neuronal PK15 recipient cells were seeded onto the SCG axons in the neurite chamber (Fig 2A). Neurons in the left soma chamber were infected with either PRV Becker, or PRV 180 that expresses a fluorescent capsid protein (mRFP-VP26), and treated with TTX (4 μM) or KCl (55 mM) from 1 hpi. We monitored the extent of virus spread to recipient PK15 cells by live-cell fluorescence microscopy, acquiring large tiled images of the entire neurite compartment every 20 min. beginning at 10 hpi (Fig 2B-C). We also performed plaque assays measuring virus titer in the neurite compartment (Fig 2D). The first fluorescent capsid protein (mRFP-VP26) expression was detected by 16 hpi in all conditions (Fig 2B), and we did not observe any differences in the kinetics of spread over time (Fig 2C). TTX treatment had no effect on virus titer in the neurite compartment at 17 or 24 hpi, and KCl treatment caused an ~1 log reduction in neurite compartment titer at 17 hpi.

**Fig 2.**
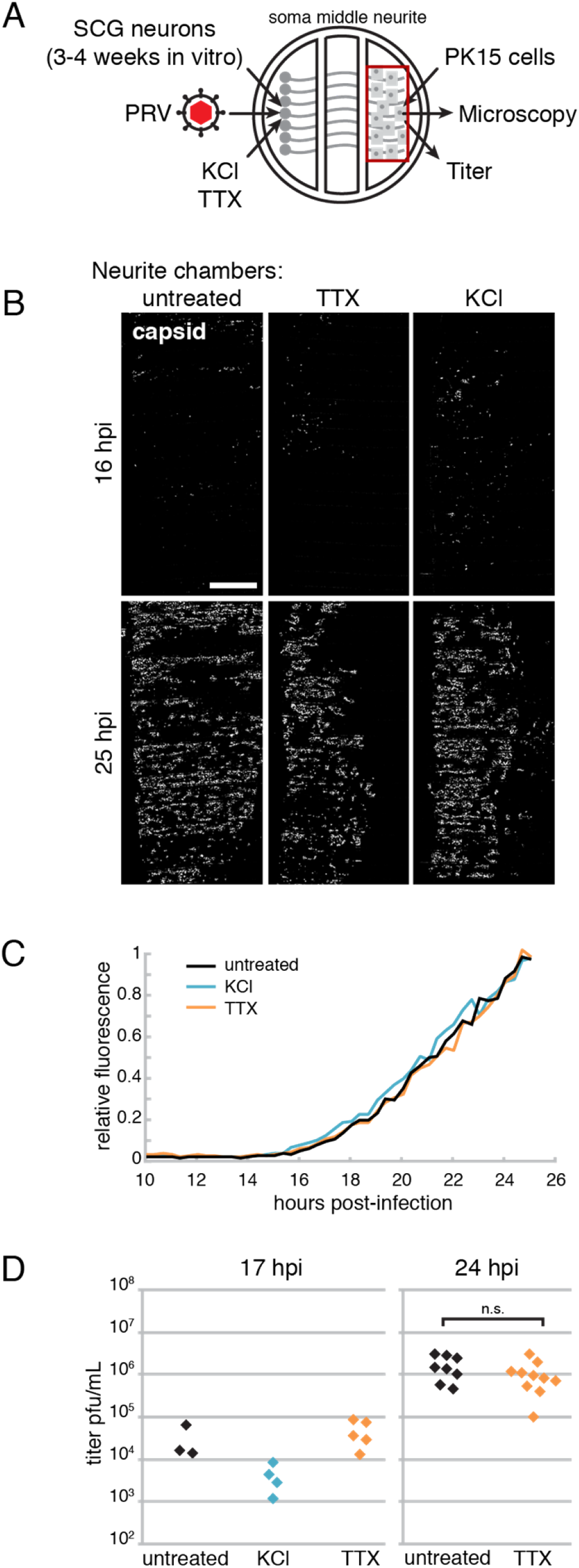
Neuronal activity does not correlate with PRV spread from neurons to non-neuronal cells. (A) Schematic of modified Campenot tri-chamber. SCG neurons were seeded in the left soma compartment and extended axons under the chamber walls to the right neurite compartment. A detector cell monolayer (PK15) was then added to the neurite compartment. PRV infection was initiated in the soma compartment, and neurons were treated with 4μM TTX, or pulses of 55mM KCl, or untreated, as indicated. (B) Tiled image of neurite chambers (area indicated by red box in panel A). Scale bar represents 1mm. (C) Relative mRFP-VP26 capsid fluorescence of entire neurite chambers over time. (D) Virus titer measured in neurite chambers by serial dilution plaque assay.

### Neuronal activity does not promote PRV spread from neuron to neuron

To determine whether neuronal activity affects spread of PRV from neuron to neuron, we prepared tri-chamber cultures with SCG neurons as recipient cells. We first cultured SCG neurons in the left soma compartment of Campenot tri-chambers for 3 weeks, to allow axon extension into the right neurite compartment. We then seeded additional freshly-prepared SCG neurons onto the axons in the neurite compartment and maintained these cultures for an additional 1 week (Fig 3A). The Banker laboratory previously described such “heterochronic cultures” (i.e. co-culture of neurons of different ages) and observed synapses forming between hippocampal neurons within 1-3 days [24]. It is well established that dissociated SCG neurons form functional adrenergic and cholinergic synapses in vitro (discussed in [25,26]), and synaptogenesis can be detected within 6-7 days in vitro [27–29].

**Fig 3.**
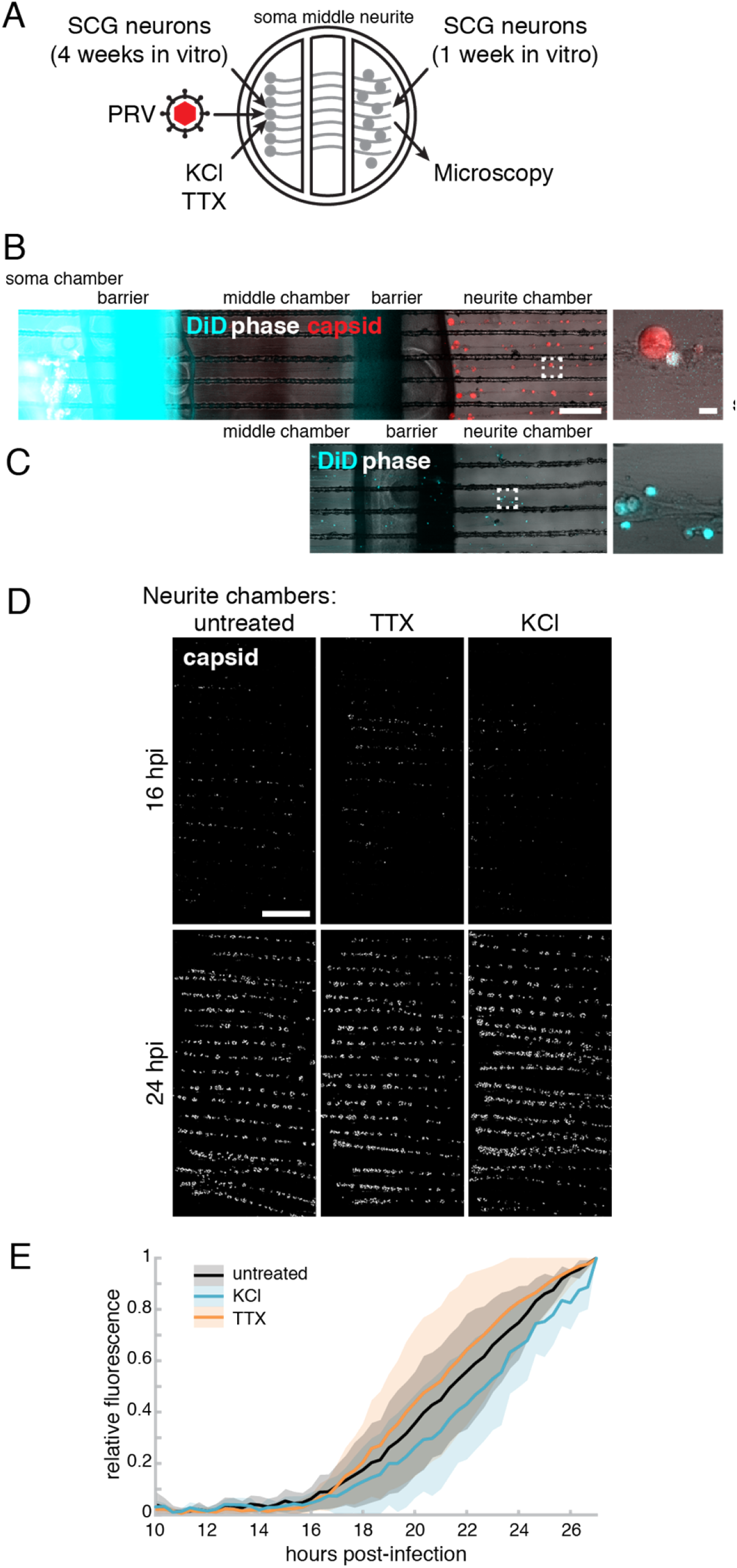
Neuronal activity does not correlate with PRV spread from neuron to neuron. (A) Schematic of modified Campenot tri-chamber. SCG neurons were seeded in the left soma compartment and extended axons under the chamber walls to the right neurite compartment. Recipient SCG neurons were then added to the neurite compartment. PRV infection was initiated in the soma compartment, and neurons were treated with 4μM TTX, or pulses of 55mM KCl, or untreated, as indicated. (B) Recipient neurons in the right neurite chamber do not penetrate the left soma compartment. Lipophilic fluorescent dye, DiD (blue), was added to the soma chamber, and chambered cultures were imaged by live-cell fluorescence microscopy. Out of many chambers, only a single DiD-labeled cell body was detected (zoom). Scale bars represent 0.5 mm (left) and 30 μm (zoom). (C) A defective leaky chamber demonstrates extensive DiD-labeling of cells, demonstrating that these imaging parameters can readily detect DiD-positive cells in the neurite chamber. (D) Tiled image of neurite chambers (area indicated by red box in Figure 2A). Scale bar represents 1mm. (E) Mean relative mRFP-VP26 capsid fluorescence of entire neurite chambers over time. Mean of n=8 independent experiments. Shaded area represents standard deviation. No significant difference (p>0.05) at all time points.

To ensure that newly-added recipient SCG neurons cannot extend axons though the chamber barriers into the left soma chamber within 1 week, we added the lipophilic far-red fluorescent dye DiD to the left soma chamber (Fig 3B-C). Any recipient SCG neurons that had extended axons into the soma chamber would become labeled by the fluorescent dye. We acquired large tiled images of entire tri-chamber cultures by fluorescence microscopy, and observed only one instance of a single DiD-labeled recipient neuron in the neurite chamber (Fig 3B), whereas a defective leaky chamber had a much greater amount of DiD labeling, demonstrating that our imaging parameters can readily detect DiD-positive cells (this defective chamber was not used for any subsequent experiments) (Fig 3C). Neurons in the left soma chamber were infected with PRV 180, expressing a fluorescent capsid protein (mRFP-VP26), then TTX (4 μM) was added beginning at 5 hpi, or KCl (55 mM) was perfused in pulses beginning at 6 hpi (Fig 3A). The extent of virus spread to recipient neurons in the neurite chamber was monitored by fluorescence microscopy, acquiring large tiled images of the entire neurite compartment every 20 min. beginning at 10 hpi. In all conditions, the first fluorescent capsid protein (mRFP-VP26) expression was detected by 16 hpi (Fig 3D), and we did not observe any significant (p>0.05) differences in the kinetics of spread over time (Fig 3E).

### Neuronal activity does not promote PRV spread from neurons: Optogenetics Approach

Because KCl treatment causes excitotoxicity in neurons and has been previously reported to reduce HSV-1 replication [30], we next pursued an optogenetics approach to modulate neuronal firing activity. Channelrhodopsin 2 (ChR2) is a light-gated ion channel originally isolated from algae, which has become a popular tool to control the activity of neurons using light [31]. To validate this method, we first plated SCG neurons on a multi-electrode array, transduced these neurons with an AAV vector to express ChR2-mCherry (Fig 4A-B), and performed extracellular recordings of their spontaneous and light-evoked spiking activity (Fig 4C). Greater than 80% of neuronal cell bodies expressed ChR2-mCherry red fluorescence (Fig 4A-B). Without blue light stimulus, neurons exhibited an average spontaneous spiking frequency of 4 Hz. Neurons were then exposed to blue light pulsed at approximately 12 Hz. With blue light stimulation, neuron activity was synchronized to the pulsed blue light, with a measured average spiking frequency of 11 Hz. Light stimulation also increased peak-to-peak spike amplitude from an average of 70 μV to 108 μV (Fig 4C-D).

**Fig 4.**
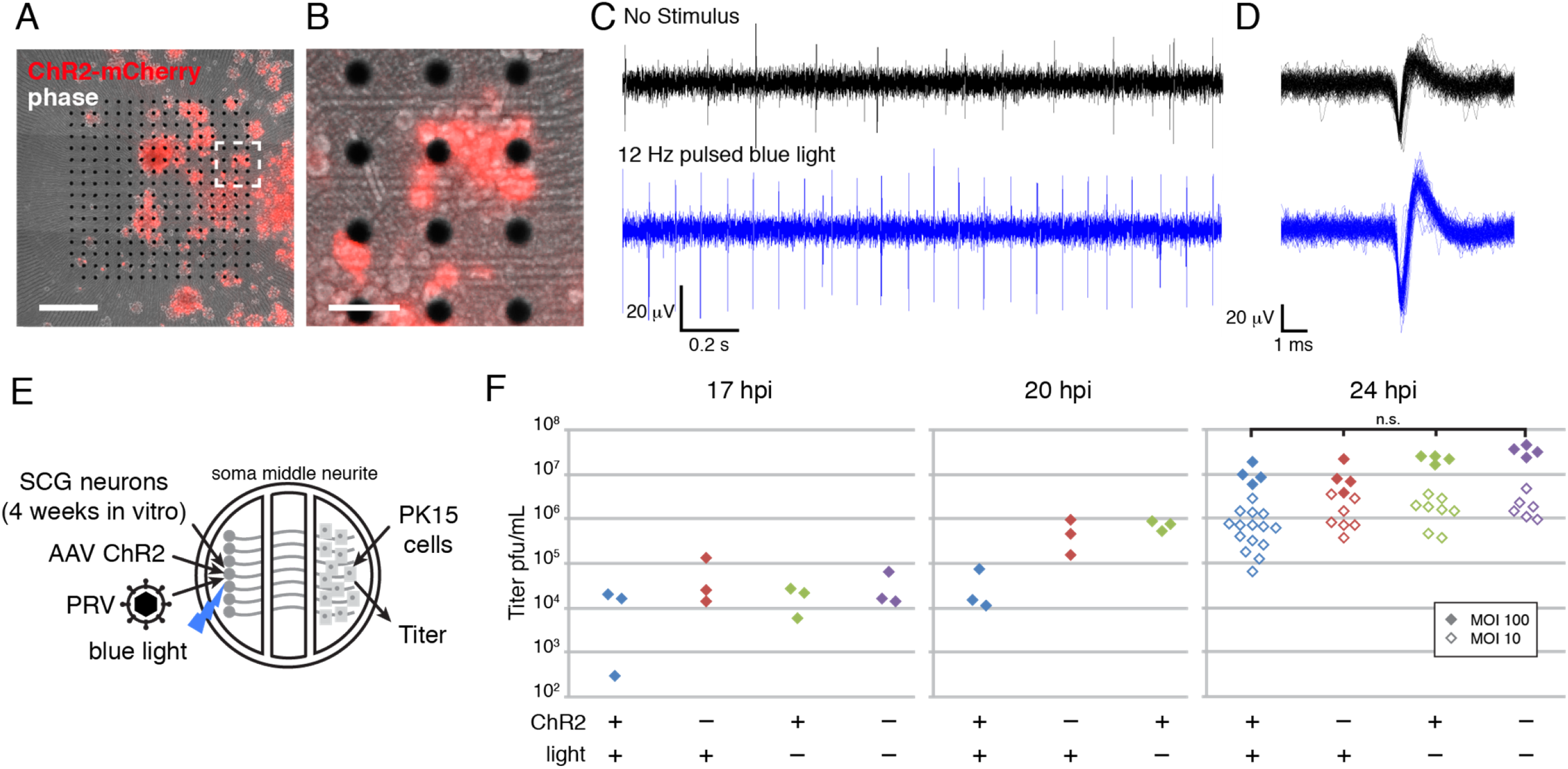
Neuronal activity does not correlate with PRV spread from neurons: optogenetics approach. (A-B) SCG neurons were seeded onto a multi-electrode array, transduced with an AAV vector expressing ChR2-mCherry. (A) Scale bar is 500μm. (B) Scale bar is 100μm. (C) Extracellular electrical recordings of SCG neurons exhibiting spontaneous spiking activity (no stimulus), or evoked activity synchronized to 12 Hz blue light pulses. (D) Superimposed spikes (n=100) demonstrating increases in spike peak-to-peak amplitude with optogenetic stimulation. (E) Schematic of modified Campenot tri-chamber. SCG neurons were seeded in the left soma compartment and extended axons under the chamber walls to the right neurite compartment. A detector cell monolayer (PK15) was then added to the neurite compartment. SCGs were transduced with a ChR2 AAV and exposed to pulsed blue light, as indicated. PRV infection was initiated in the soma compartment at a multiplicity of infection of 10 or 100, as indicated. (F) Virus titer measured in neurite chambers by serial dilution plaque assay. No significant difference (p>0.05) at 24 hpi.

To determine whether optogentically-controlled neuronal activity affects spread of PRV, we prepared tri-chambers with SCG neurons in the left soma chamber, and transduced with an AAV vector to express ChR2-mCherry. We screened neurons by fluorescence microscopy to ensure ChR2-mCherry expression in >80% of cells. At 4 weeks in vitro, we seeded PK15 recipient cells in the right neurite compartment, infected with PRV Becker, and pulsed with blue light beginning at 1 hpi (Fig 4E). In the experimental condition, where neurons were transduced with ChR2 and pulsed with blue light, neurite compartment titers were slightly less than controls at 17 and 20 hpi, but not significantly different by 24 hpi (p>0.05) (Fig 4F).

### PRV particle exocytosis from neuronal cell bodies occurs as early as 5 hours post-infection

To directly measure PRV egress from infected cells, we previously developed a live-cell fluorescence microscopy assay of virus particle exocytosis [14,15]. We genetically fused pHluorin, a pH-sensitive variant of EGFP, to the first extracellular loop of viral glycoprotein M (gM-pHluorin). When gM-pHluorin is incorporated into virus particles and surrounding secretory vesicle membrane during secondary envelopment, pHluorin fluorescence is strongly quenched in the acidic lumen of the secretory organelle. Upon exocytosis, the pHluorin moiety is exposed to the higher extracellular pH and becomes brightly fluorescent, revealing the location and moment of virus particle egress [14,15]. We infected primary SCG cultures with PRV 483, expressing gM-pHluorin and fluorescent capsid protein (mRFP-VP26), and imaged by Total Internal Reflection Fluorescence (TIRF) microscopy. In agreement with previous work in non-neuronal cells, we detected exocytosis of individual virus particles based on the sharp increase in gM-pHluorin fluorescence. We detected exocytosis of both virions, categorized based on the colocalization of gM-pHluorin and mRFP-VP26 capsid fluorescence, and light particles, categorized based on the absence of mRFP-VP26 (Fig 5A-B). Previously, we observed that egress of PRV virions occurs as early as 4.5-5 hpi in non-neuronal PK15 cells [14]. Here we observed PRV particles undergoing exocytosis from the cell body of SCG neurons as early as 5-6 hpi, much earlier than the previously-reported increases in neuronal activity and increases in intracellular Ca^2+^ at 8-10 hpi [10,22].

**Fig 5.**
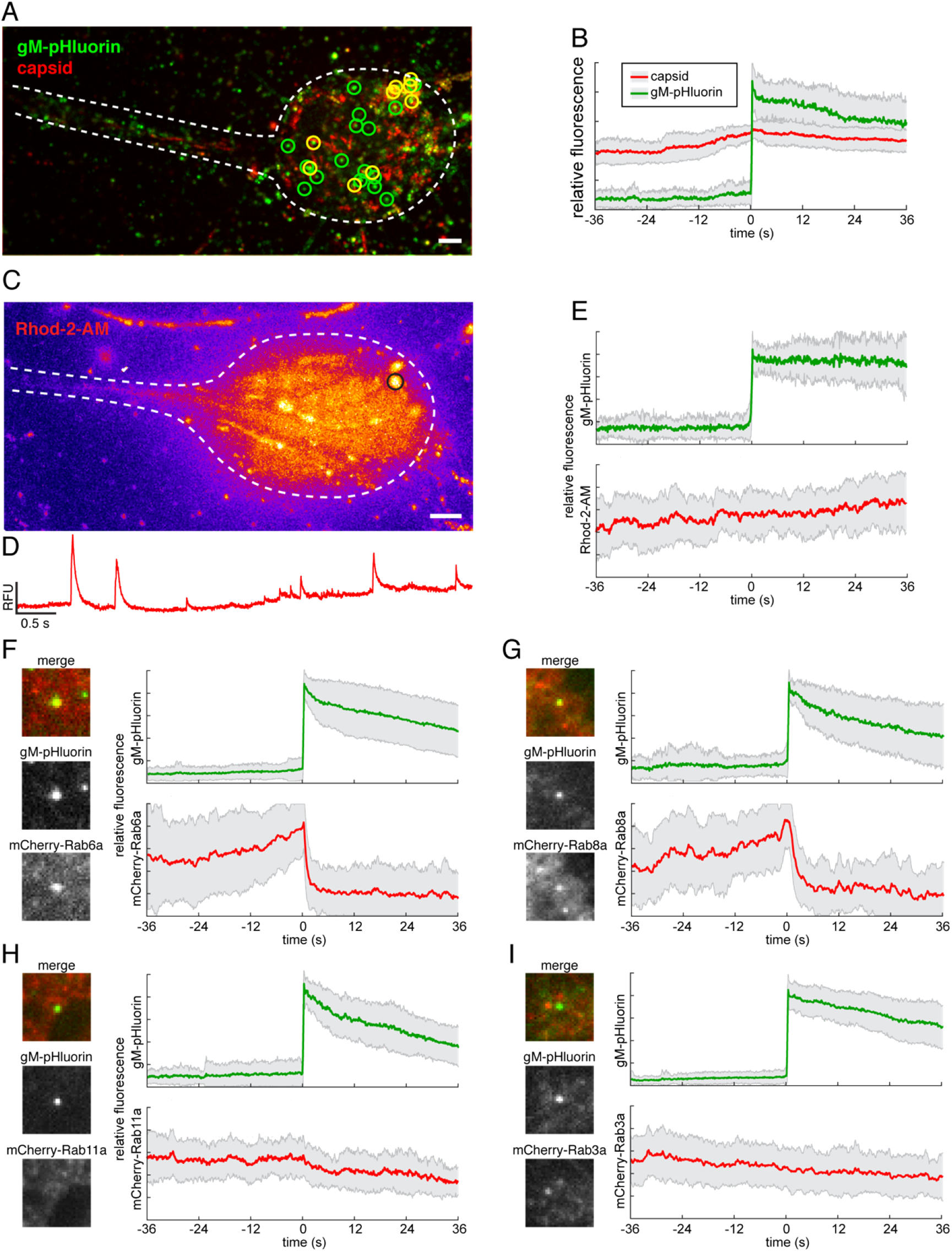
Live-cell fluorescence microscopy of virus exocytosis in primary SCG neurons. (A) Virus particle exocytosis in neurons occurs as early as 5 hpi. Image is a maximum difference projection over a 5.3 min time course. Exocytosis of gM-pHluorin particles that do not contain capsids (green circles) and particles containing both gM-pHluorin and capsids (yellow circles) are indicated. Scale bar represents 4 μm. (B) Ensemble average of gM-pHluorin fluorescence (green line) and mRFP capsid (red line) over 26 exocytosis events. Shaded area represents standard deviation. (C) Neurons were infected with PRV expressing gM-pHluorin, loaded with Ca^2+^ sensitive fluorescent dye Rhod-2-AM, and imaged beginning at 5 hpi. Image is a maximum intensity projection of Rhod-2-AM fluorescence over a 6 min time course. Scale bar represents 4 μm. (D) Rhod-2-AM fluorescence intensity at the region of interest indicated in panel C (black circle) over 6 min time course. (E) Virus particle exocytosis in neurons is not associated with local Ca^2+^ transients. Ensemble average of gM-pHluorin (top, green line) and Rhod-2-AM (bottom, red line) relative fluorescence over 27 exocytosis events. Shaded area represents standard deviation. (F-G) Neurons were transduced to express mCherry-tagged Rab proteins, infected with PRV expressing gM-pHluorin, and imaged beginning at 5 hr after PRV infection. Images show representative exocytosis events at the moment of exocytosis (time=0). Each image is 5 μm square. Line plots are ensemble averages of gM-pHluorin (top, green line) and indicated Rab protein (bottom, red line) relative fluorescence. Shaded area represents standard deviation. (F) Rab6a is associated with virus particle exocytosis in neurons. Data represent 72 exocytosis events. (G) Rab8a is associated with virus particle exocytosis in neurons. Data represent 30 exocytosis events. (H) Rab11a is not associated with virus particle exocytosis in neurons. Data represent 41 exocytosis events. (I) Rab3a is not associated with virus particle exocytosis in neurons. Data represent 64 exocytosis events.

### PRV particle exocytosis from neuronal cell bodies is not correlated with local Ca^2+^ dynamics

In addition to global membrane depolarization and Ca^2+^ transients associated with neuronal firing activity, neurons and other excitable cell types, including SCG neurons in particular [32], also exhibit smaller localized Ca^2+^ transients, variously described as “Ca^2+^ sparks”, “Ca^2+^ glows”, or “syntillas”. These local Ca^2+^ dynamics are highly variable in amplitude, time course, and likely result from a variety of intracellular signaling pathways [32–36]. To determine whether virus exocytosis events are correlated with local Ca^2+^ dynamics, we infected primary SCG cultures with PRV 486, expressing gM-pHluorin, loaded cells with the membrane permeant Ca^2+^-sensitive dye Rhod-2-AM, and imaged beginning at 5 hpi. Infected neurons exhibited local Ca^2+^ dynamics indicated by local changes in Rhod-2-AM fluorescence intensity (Fig 5C-D). To generalize the relationship between fluorescent signals, we measured the gM-pHluorin and Rhod-2-AM fluorescence of many exocytosis events, aligned them to a common timescale based on the moment of exocytosis, and calculated an ensemble average showing the relative fluorescence intensity over time. In this analysis, virus exocytosis events were not associated spatially or temporally with local Ca^2+^ dynamics (Fig 5E), consistent with previous studies indicating that individual local Ca^2+^ transients are not correlated with spontaneous exocytosis in uninfected neurons and chromaffin cells [34,35].

### PRV particle exocytosis from neuronal cell bodies uses constitutive secretory pathway

To characterize the viral secretory vesicle and identify cellular factors involved in PRV egress, we transduced primary SCG cultures with non-replicating adenovirus vectors expressing mCherry red fluorescent protein-tagged Rab GTPases. Approximately 18 hr after transduction, we infected with PRV 486, expressing gM-pHluorin, and imaged cell bodies 5-6 hr after PRV infection. To generalize the relationship between fluorescent signals, we measured gM-pHluorin and mCherry-Rab GTPase fluorescence of many exocytosis events, aligned them based on moment of exocytosis, and calculated an ensemble average of relative fluorescence intensity over time (Fig 5F-I). An increase in Rab fluorescence before exocytosis represents the gradual arrival of Rab-positive secretory vesicles to their sites of exocytosis. After exocytosis, Rab protein fluorescence rapidly decays, representing diffusion away from the site of exocytosis, consistent with the dynamic regulation of Rab activity and membrane binding by GTP hydrolysis (Fig 5F-I). Using this method, we found that Rab6a and Rab8a, which mediate constitutive exocytosis of post-Golgi secretory vesicles, are dynamically associated with viral exocytosis (Fig 5F-G), in agreement with our previous results in non-neuronal cells [14,15]. In contrast, we previously observed that Rab11a, which mediates trafficking of recycling endosomes, is also associated with viral secretory vesicles in non-neuronal cells; however, we show here that Rab11a does not appear to be associated with virus particle exocytosis in neurons (Fig 5H), consistent with the divergent roles of Rab11 proteins in neurons versus non-neuronal cells [19]. Finally, Rab3a, which mediates the trafficking and Ca^2+^-regulated exocytosis of synaptic vesicles in neurons, also does not appear to be associated with virus particle exocytosis (Fig 5I). Altogether, these results are consistent with PRV particles using the constitutive secretory pathway, not Ca^2+^-regulated synaptic vesicle-like secretory mechanisms, even in neurons.

## Discussion

Based on three main lines of evidence, 1) that alpha herpesviruses spread in a synaptic circuit-specific manner in the mammalian nervous system; 2) that alpha herpesvirus proteins and particles associate with synaptic vesicle markers; and, 3) that alpha herpesviruses cause an elevated rate of neuronal activity *in vitro* and *in vivo*, it has long been suspected that alpha herpesviruses take advantage of synaptic vesicle-like regulated secretory mechanisms in neurons. Here we present four pieces of evidence that contradict this view. Using Campenot tri-chambers to measure transneuronal virus spread, and tetrodotoxin to silence neuronal activity, we observed no inhibition of virus spread from primary neurons. Using KCl or optogenetics to elevate neuronal activity, we also show no increase in virus spread. Using a live-cell fluorescence microscopy method to directly assay virus egress from infected neurons, we observed no association between virus particle exocytosis and intracellular Ca^2+^ signaling. Finally, we observed virus particle exocytosis occurs in association with constitutive secretory Rab GTPases, Rab6a and Rab8a, not Rab proteins that are associated with the Ca^2+^-regulated secretory pathway in neurons, Rab3a and Rab11a. Therefore, we conclude that alpha herpesvirus egress and spread is independent of neuronal activity and Ca^2+^ signaling because virus particle exocytosis uses constitutive secretory mechanisms in neurons.

### Clinical Implications

These observations have several important clinical implications. Our findings suggest the altered electrophysiology and elevated activity is an epiphenomenon of alpha herpesvirus infection in neurons and is not required for productive assembly and egress of progeny virions. Therefore, medical interventions treating sensory neuropathic symptoms or targeting regulated secretory mechanisms in neurons (e.g. neurotoxins) are unlikely to inhibit the spread of alpha herpesviruses in the nervous system. However, therapeutic strategies that target cellular factors common to both regulated and constitutive secretory pathways might inhibit virus egress and spread. For example, botulinum neurotoxin A (Botox) cleaves the SNARE protein SNAP-25, which is required for both Ca^2+^-regulated exocytosis of synaptic vesicles (hence a neurotoxin) as well as intracellular trafficking and constitutive exocytosis in neurons and non-neuronal cells [37]. Botox has been tested as a potential treatment to prevent recurrence of herpes labialis, but aside from limited case reports from off-label use [38], the results from clinical trials have not been published.

### Future Directions

There are several important limitations of our approaches. Foremost, while SCG neurons do form functional synapses *in vitro*, measuring transneuronal virus spread through the Campenot tri-chamber cannot distinguish between spread via these neuronal synapses and spread via other non-synaptic cell-cell contacts. It is therefore possible that alpha herpesviruses do spread via neuronal synapses in an activity-dependent manner, but this was not detected in our assays because the virus also spreads efficiently via non-synaptic cell-cell contacts in an activity-independent manner. It is important to note that in this study, using our live-cell fluorescence microscopy method, we only observed virus particle exocytosis from neuron cell bodies. It is therefore possible that alpha herpesviruses use the constitutive secretory pathway for egress from the cell body and other non-synaptic egress sites, but use Ca^2+^-regulated secretory mechanisms for egress specifically at neuronal synapses. To resolve these remaining ambiguities, in future studies it will be necessary to directly observe virus egress and spread specifically at neuronal synapses to distinguish the molecular and cellular mechanisms of synaptic versus non-synaptic cell-cell spread.

## Materials and Methods

### Cells

PK15 cells (ATCC CCL-33) were cultured in Dulbecco’s Modified Eagle Medium (DMEM) supplemented with 10% fetal bovine serum (FBS) and 1% penicillin/streptomycin, at 37°C and 5% CO_2_. Primary embryonic rat superior cervical ganglion (SCG) neurons were isolated and cultured as previously described [23]. To measure anterograde spread, neurons were cultured in modified Campenot tri-chambers, as previously described [23]. For TIRF microscopy, SCG neurons were cultured on 35mm glass-bottom tissue culture dishes (Mattek), which were treated by corona discharge (Electro-Technic Products) and sequentially coating with poly-DL-ornithine and laminin. For extracellular electrode recording, SCG neurons were cultured on a multielectrode array (Multichannel Systems), which was prepared by sequentially coating with polyethylenimine (PEI) and laminin. In all cases, neurons were cultured in Neurobasal medium supplemented with B-27 (ThermoFisher), glutamine, penicillin/streptomycin, and nerve growth factor (NGF), as previously described [23], for 3-4 weeks to achieve a fully-differentiated and polarized neuronal phenotype.

### Ethics Statement

Animals were euthanized by carbon dioxide inhalation, as recommended by the American Veterinary Medical Association (AVMA). All animal work was approved by the Princeton University Institutional Animal Care and Use Committee (protocol #1947), in accordance with applicable policies and laws, including the Animal Welfare Act (AWA), the Public Health Service Policy on Humane Care and Use of Laboratory Animals, the Principles for the Utilization and Care of Vertebrate Animals Used in Testing, Research and Training, and the Health Research Extension Act of 1985.

### Viruses

All recombinant viruses are derivatives of PRV Becker, a wild-type laboratory strain [39]. PRV 180, expressing an mRFP-VP26 capsid protein fusion, was previously described [40]. PRV 468, expressing mRFP-VP26 and the fluorescent protein Ca^2+^ sensor GCaMP3, was previously described [11]. PRV 486, expressing mRFP-VP26 and gM-pHluorin, was previously described [14]. SCG neurons were infected with PRV recombinants at a multiplicity of infection (MOI) of approximately 5 pfu/cell, unless otherwise in the figure or figure legend. Adenovirus vectors expressing mCherry-tagged Rab3a, Rab6a, Rab8a, and Rab11a were previously described [14]. SCG neurons were transduced with adenovirus vectors approximately 18 hours before co-infection with PRV. An adeno-associated virus (AAV) vector expressing channelrhodopsin2 (ChR2) fused to mCherry (AAV2/1.CAG.hChR2(H134R)-mCherry.WPRE.SV40) was obtained from the U. Pennsylvania Vector Core. SCG cultures were inoculated with 10^7^ transducing units of AAV and screened by fluorescence microscopy for >80% transduction efficiency.

### Optogenetics and Extracellular Electrode Recording

SCG neurons transduced with AAV vectors expressing ChR2 were exposed using a blue LED light box with an excitation spectrum of approximately 457/46 nm (Clare Chemical Research, Denver, CO, USA) [41], driven by an Arduino Uno Rev3 microcontroller programmed to deliver light pulses at approximately 12 Hz (see S1 Supporting Materials and Methods). Extracellular recordings of spontaneous and light-evoked spiking activity were digitized at 10 kHz and stored for offline analysis, as previously described [42]. The extracellular recordings were composed of fast negative voltage spikes from individual neurons, superimposed on a slowly fluctuating local field potential generated by the average activity of many nearby neurons. Recordings were processed by subtracting electronic noise recorded on a reference electrode and 200 Hz high-pass filtered to remove local field potentials, spikes were identified using a threshold of −30 μV, and average spike frequency and peak-to-peak amplitude were measured using MC_Rack software (Multichannel Systems).

### Fluorescence Microscopy and Image Processing

Tiled images of entire neurite compartments in tri-chambers were acquired using an automated epifluorescence microscope equipped with a 37°C and 5% CO_2_ environmental chamber for long-term imaging of live cells, as previously described [43]. To measure the total fluorescence in chambered neuronal cultures, we performed background subtraction and feature selection using median filter background subtraction and granulometric filtering in Fiji/ImageJ v.1.52b software, as previously described [44]. Total Internal Reflection Fluorescence (TIRF) microscopy was performed using a Nikon N-STORM fluorescence microscope, as previously described [14,15]. Images were prepared for publication using the following functions and plugins in Fiji/ImageJ: adjust brightness and contrast, Z project (to make maximum intensity projections). We calculated maximum difference projections in Fiji/ImageJ as follows: Image values at time n were subtracted from values at time n+5 to identify pixels where fluorescence intensity increases, and the resulting image stacks were then processed by maximum intensity projection to highlight areas where fluorescence intensity increases the most.

## Acknowledgements

Thank you to members of the Lynn Enquist, Michael Berry, Joshua Shaevitz, Sam Wang, and Esteban Engel laboratories at Princeton for technical advice and assistance. Thanks to Gary Laevsky at the Dept. of Molecular Biology Confocal Microscopy Facility for microscopy assistance. This research was supported by National Institutes of Health grants R01 NS060699 (LWE) and K22 AI123159 (IBH).

## Supporting Information

### S1 Supporting Materials and Methods

#### Arduino Controller for Optogenetics Light Stimulation

An Arduino controller (Arduino Uno R3, Arduino, Milan, Italy) was built and programmed to control of a relay switch (PowerSSR Tail, powerswitchtail.com, Honolulu, HI), timing the on/off cycle of a transilluminator (Clare Chemical Research, Dolores, CO) to drive channelrhodopsin-2 activity in long-term optogenetics experiments inside a cell culture incubator.

Two dial potentiometers control the on rate and off rate at frequencies up to 30 Hz. Due to response delay in the lightbox, the reported on and off times should be treated as a reference only (reported on time is an overestimate and off time is an underestimate). Measurements should be made with a photosensor if more precise on/off rate is sought.

Below is the code for the Arduino controller that is designed to interact with circuitry illustrated in Figure S1. The code is written in the Arduino programming language, which is derivative of C/C++.

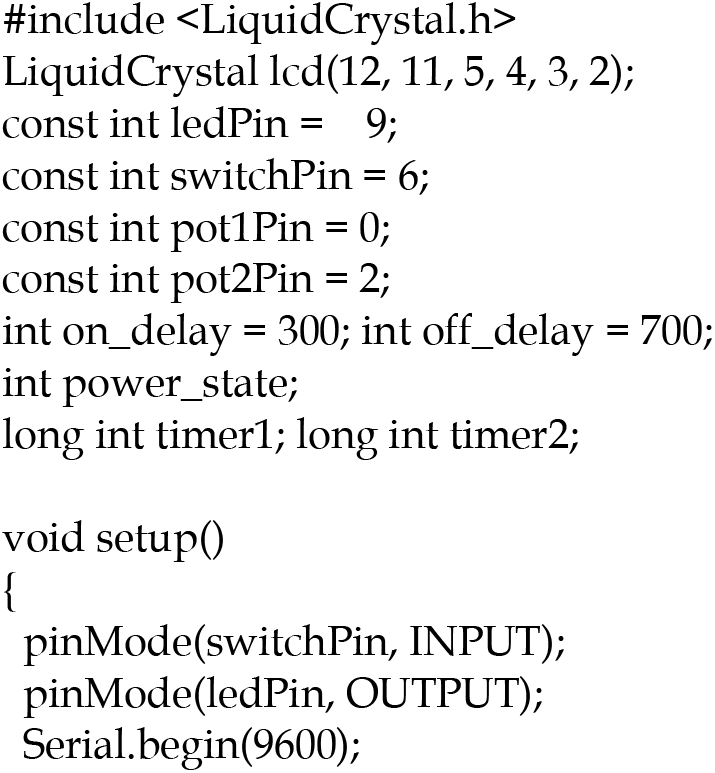
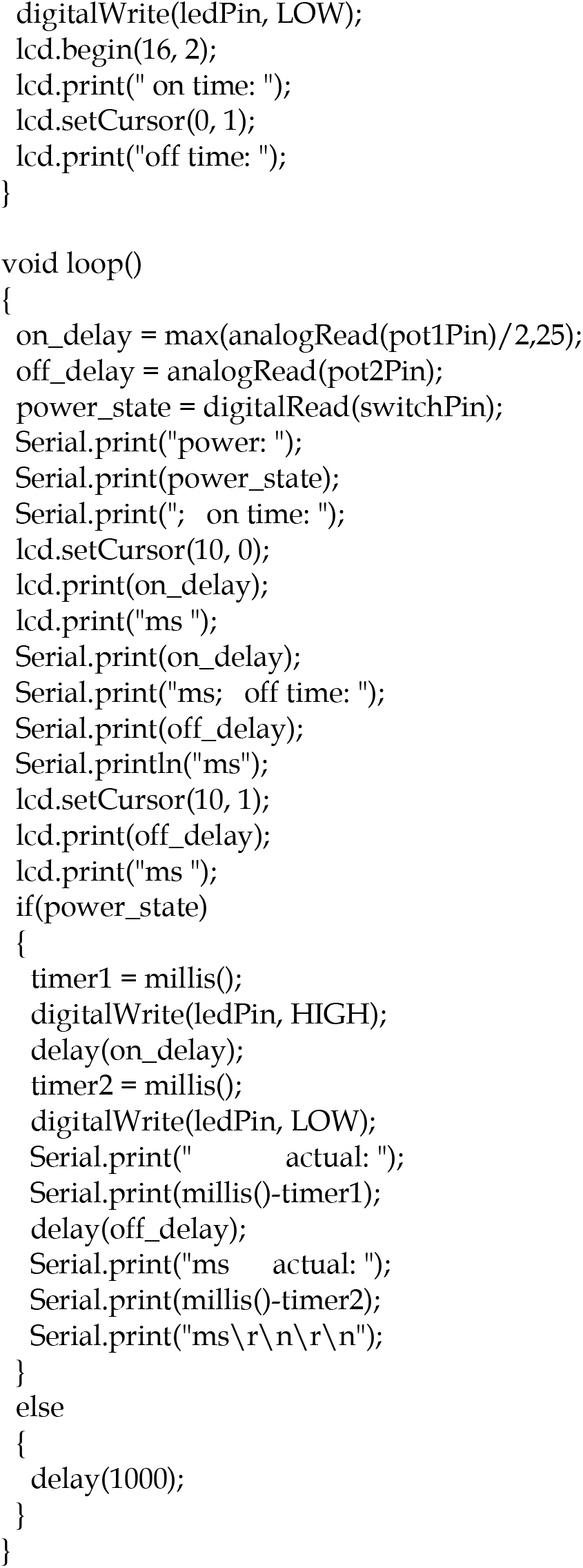

**S1 Figure.**
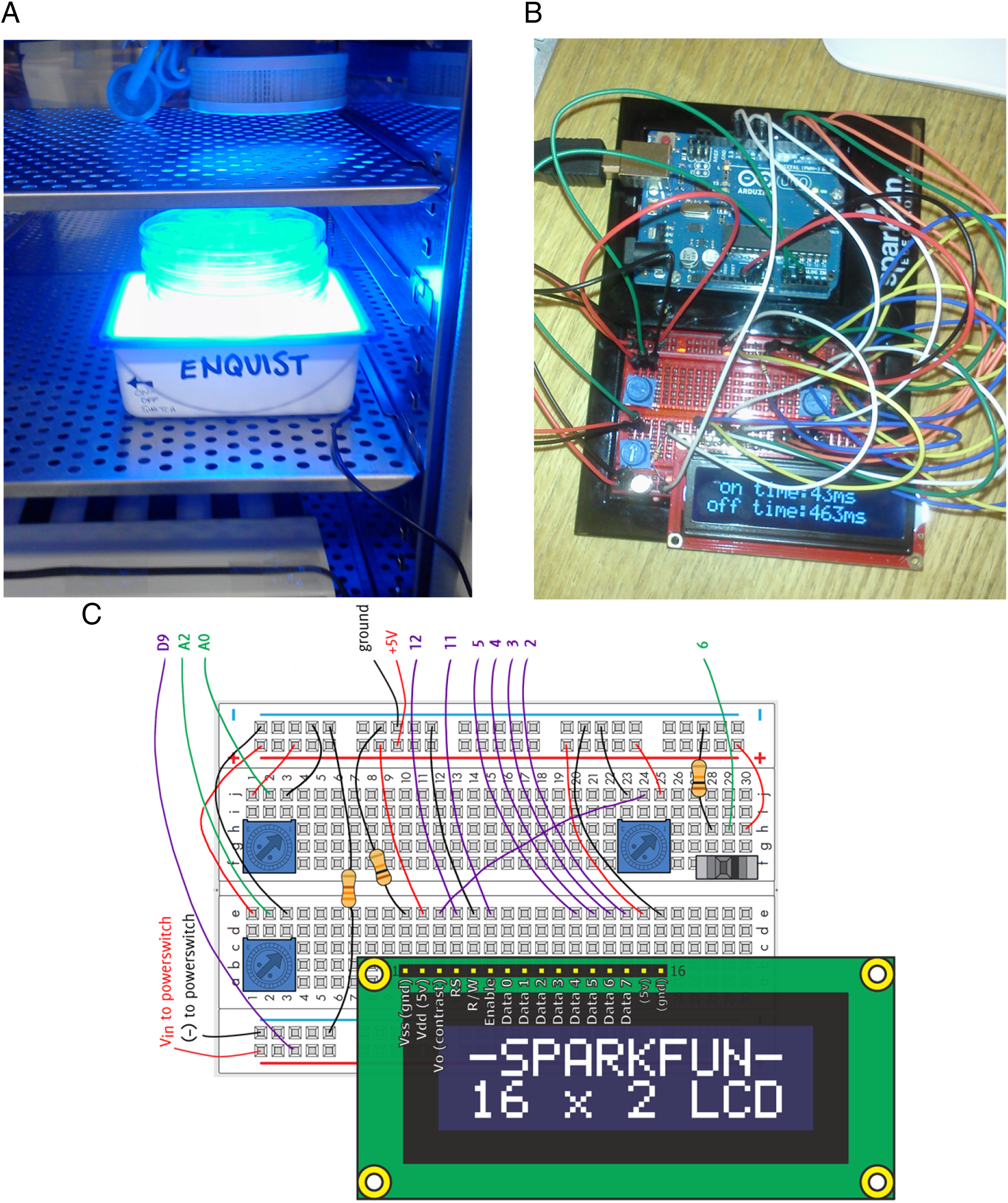
Arduino microcontroller circuitry for optogenetics light stimulation. (A) Photograph of blue light transilluminator with cell culture dishes installed in a cell culture incubator. (B) Photograph of Arduino microcontroller with wiring (arbitrary wire colors). (C) Wiring diagram for Arduino controller. Inputs and outputs are color-coded green and purple, respectively. Potentiometers connected to inputs A0 and A2 control the relay switch on and off times, respectively. The third potentiometer controls the contrast on the LCD display, where an estimate of on and off times is displayed. The switch toggles the system on and off.

